# A novel recombinant BCG vaccine using a mycobacteriophage promoter shows improved protection against tuberculosis

**DOI:** 10.64898/2026.09.04.749280

**Authors:** Yumiko Tsukamoto, Yusuke Tsujimura, Tetsu Mukai, Toshiki Tamura, Yumi Maeda, Shota Torigoe, Hiroyuki Saiga, Masamitsu N. Asaka, Takeshi Komine, Yuji Miyamoto, Makoto Nakaya, Masahiko Makino, Manabu Ato

**Author notes:** Corresponding Author: Yumiko Tsukamoto E-mail address, Address: 4-2-1 Aobacho, Higashimurayama-shi, Tokyo, Japan.

## Abstract

The limited efficacy of conventional bacillus Calmette–Guérin (BCG) vaccination against adult pulmonary tuberculosis highlights the need for improved vaccine strategies. Genome integration can overcome the plasmid instability of recombinant BCG, but we hypothesized that this approach would be constrained by reduced transcriptional activity of conventional promoters when present as a single genomic copy. Here we identified P79, a mycobacteriophage-derived promoter that sustains high-level expression from a single genomic locus and used it to construct a genome-integrated recombinant BCG, termed LRC-BCG. LRC-BCG secretes the HSP70-MMPII fusion antigen flanked by the PEST sequences, maintained stable antigen expression over serial passages, and induced strong activation of macrophages and dendritic cells in vitro. *In vivo,* LRC-BCG inhibited *Mycobacterium tuberculosis* multiplication more effectively than conventional BCG, even at low doses. These findings demonstrate the feasibility of using P79 for stable genome-integrated recombinant BCG and support LRC-BCG as a promising tuberculosis vaccine candidate.

## Introduction

*Mycobacterium tuberculosis* (MTB) is the causative agent of tuberculosis (TB), a chronic infectious disease that poses a major global health burden. In 2024, TB caused an estimated 1.23 million deaths worldwide, making it the leading cause of death from a single infectious agent globally, surpassing COVID-19^1^. Although effective antimicrobial therapies are available, the emergence and spread of drug-resistant TB, particularly multidrug-resistant strains, continue to pose a serious challenge for TB control. These circumstances underscore the persistent and urgent need for effective TB vaccines.

*Mycobacterium bovis* bacillus Calmette–Guérin (BCG) is a live-attenuated vaccine developed in the early 20th century and has been used for more than a century as the only licensed vaccine against TB^2–4^. The World Health Organization recommends BCG vaccination in countries with a high incidence of TB^5^, and in countries with routine BCG vaccination for all infants, BCG coverage was approximately 88% in 2022^1^. Numerous studies have demonstrated that BCG has substantial efficacy in preventing TB in children and infants, especially in cases of meningeal and disseminated tuberculosis^6–8^. However, its effectiveness against TB, especially pulmonary TB, in the adult population remains comparatively low^9–11^. This fundamental limitation of conventional BCG has driven extensive global efforts to develop improved TB vaccines^12,13^.

We previously identified the Major Membrane Protein-II (MMPII) antigen as an immunodominant antigen in *Mycobacterium leprae*^14,15^. MMPII is encoded by the *bfrA* gene and functions as a ligand for Toll-like receptor 2 (TLR2), conferring potent immunostimulatory properties through the activation of antigen-presenting cells (APCs), such as dendritic cells (DCs) and macrophages. Recombinant BCG strains expressing MMPII have been shown to enhance the activation of human APCs compared to parental BCG^15,16^, highlighting the potential of MMPII as a component of recombinant BCG vaccines.

Although MMPII was initially identified and characterized in *M. leprae*, it has also been found to be expressed in MTB and BCG. Importantly, the MMPII protein expressed by MTB and BCG was identical at the amino acid level but shared 90.6% sequence homology with its *M. leprae* counterpart. MMPII derived from MTB exhibited stronger activation of DCs and macrophages than MMPII from *M. leprae*^17^.

Therefore, we utilized the overexpression of MTB-derived MMPII to develop an improved recombinant BCG vaccine against TB^18–21^.

To further enhance antigen processing and immune activation by MTB-derived MMPII, we engineered a recombinant BCG that incorporates the *ureC* gene deletion to facilitate phagolysosomal maturation along with the secretion of an HSP70-MMPII fusion protein flanked by PEST sequences to promote efficient antigen processing and presentation. The *ureC* gene-depleted BCG (BCG-ΔUT), induced the acidification of the phagosome due to a lack of urease activity, and facilitating phagolysosome formation^22,23^. HSP70 is a chaperone protein that facilitates immune activation, and the HSP70-fusion protein is secreted by BCG^24–26^. The PEST sequence is a peptide characterized by four amino acids (P, E, S, and T)^27,28^. The PEST sequence is reported to induce the processing of various proteins in a manner dependent on host cellular components, such as the proteasome, thereby promoting intracellular antigen degradation^27–29^. We have demonstrated that recombinant BCG lacking the *ureC* gene and secreting an HSP70-MMPII fusion protein flanked by PEST sequences can activate APCs, and that vaccination with this recombinant BCG inhibited the multiplication of MTB in the lungs^18,20,21^.

This recombinant BCG was a promising candidate as a novel TB vaccine. However, we found that the recombinant plasmid was unstable after several passages of BCG. We concluded that this plasmid-based recombinant BCG is unsuitable for practical applications.

Genome integration of the antigen-coding sequence offers a straightforward solution to plasmid instability. However, this approach introduces a critical constraint: because genome-integrated constructs are present as a single copy, they require a promoter with intrinsically high transcriptional activity to achieve expression levels comparable to multicopy plasmid systems. Conventional promoters used in mycobacterial expression systems, such as Phsp60, exhibit markedly reduced activity under these single-copy conditions, and this promoter limitation has been a key barrier to the practical use of genome-integrated recombinant BCG. To overcome this bottleneck, we searched the mycobacteriophage genome for promoters capable of driving strong, single-copy expression, and identified P79, the promoter of the gp79 gene of mycobacteriophage TM4. To determine whether this promoter could support practical genome-integrated antigen expression, we generated a new recombinant BCG by integrating the promoter sequence and the fusion protein-coding sequence into the *ureC* gene locus of the host BCG and named this recombinant BCG LRC-BCG. LRC-BCG is deprived of the *ureC* gene and secretes the exogenous fusion protein. In this study, we evaluated the genetic stability of LRC-BCG across serial passages and characterized its immunostimulatory properties in macrophages and DCs *in vitro*. We also evaluated the protective efficacy of LRC-BCG against MTB infection in a murine aerosol challenge model.

## Results

### Generation of LRC-BCG and stability check

First, we sought promoters for the expression of the target antigen integrated in BCG genomic DNA (gDNA). To construct recombinant BCG strains expressing target antigens, we used plasmids based on the pMV261 backbone^18–21^. When we integrated the EGFP gene into BCG gDNA using the genome-integrating vector pMV306, we found that EGFP expression from the genome-integrated construct was markedly lower than that observed in BCG transformed with the EGFP-expressing pMV261 plasmid (Supplementary Fig. 1). As both pMV261 and pMV306 utilize the same promoter, Phsp60, the difference in expression levels can be attributed to the disparity in copy number; pMV261 exists in multiple copies within BCG cells^30^, whereas genome-integrated constructs are present as a single copy. This result confirmed that a highly active promoter would be required to achieve expression levels comparable to plasmid-based systems. We focused on the mycobacteriophage TM4 genome and identified the P79 promoter, which lies in the upstream region of the gp79 gene. To our knowledge, P79 has not previously been utilized to drive heterologous antigen expression in recombinant BCG.

As shown in Supplementary Fig. 1, the P79 promoter exhibits high transcriptional activity even when integrated into the genome. Based on these findings, we selected P79 for the construction of a genome-integrated recombinant BCG vaccine. We integrated an expression cassette composed of P79, the target antigen, and the hygromycin resistance gene cassette (Hyg^r^) into the *ureC* locus of the BCG genome (Fig. 1a and Supplementary Fig. 2). BCG-Tokyo 172 (BCG-Tokyo) was used as the host BCG. Subsequently, we removed Hyg^r^ by resolvase. The resultant recombinant BCG, LRC-BCG, is antibiotic resistance gene-free, deprived of the *ureC* gene, and expresses the target gene under the control of the P79 promoter.

**Fig. 1.**
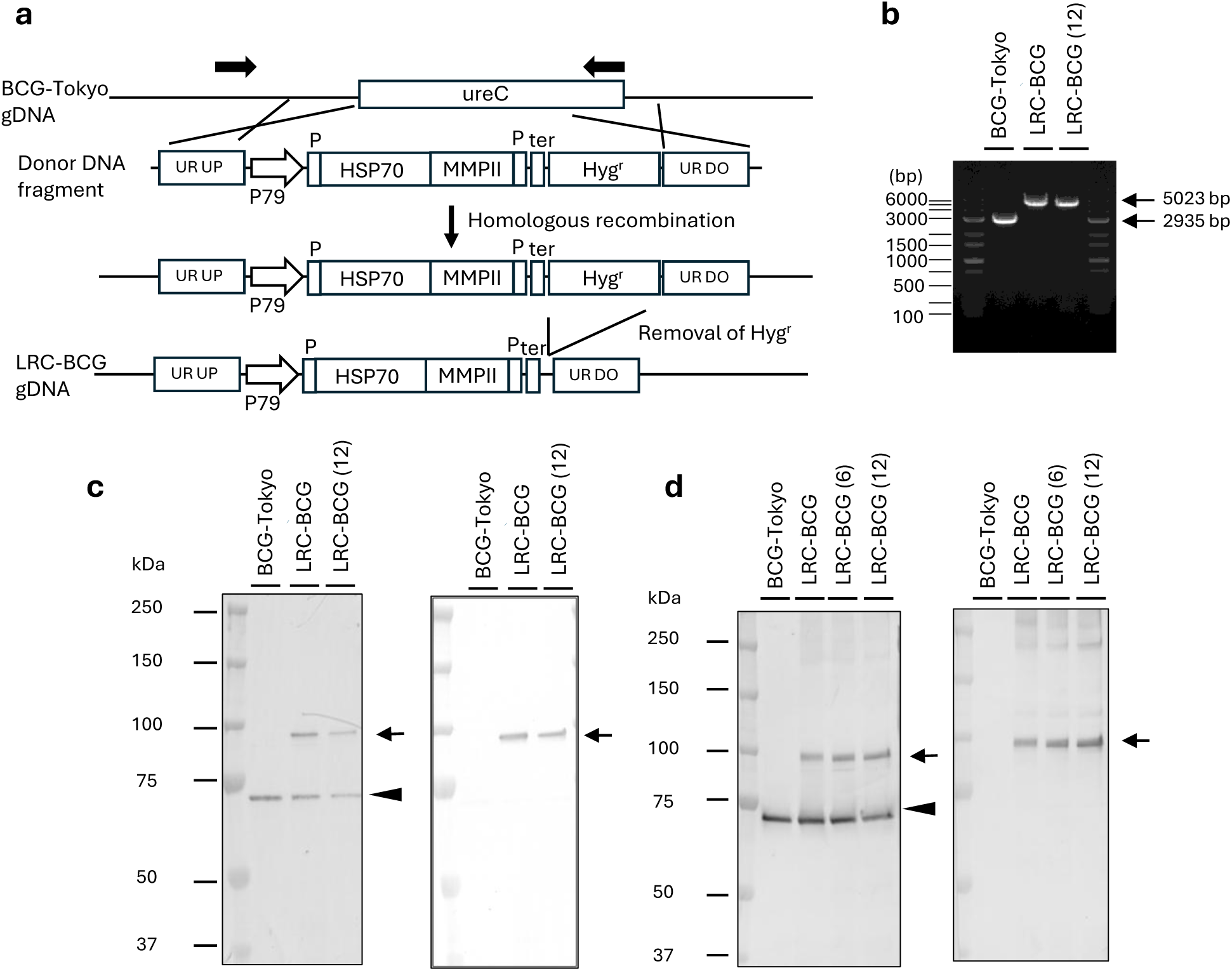
Generation of LRC-BCG. **a.** Schematic diagram of LRC-BCG generation. P and ter indicate PEST sequence and transcription terminator. UR UP and UR DO indicate the upstream and downstream regions of the *ureC* gene respectively (see Methods). The P79 promoter and PEST-HSP70-MMPII-PEST fusion gene, along with a hygromycin-resistance gene cassette (Hyg^r^), were integrated into the *ureC* gene locus of BCG-Tokyo via homologous recombination. Hyg^r^ was removed by resolvase, and the resultant LRC-BCG is antibiotic resistance gene-free. **b.** PCR analysis of the BCG, LRC-BCG, and 12 times-passaged LRC-BCG (LRC-BCG (12)) genome DNA. The sites of the primers are shown in (a) (black arrows). **c.** d. Western blotting of the bacterial pellet (c) or the culture supernatants (d) derived from BCG, LRC-BCG, 6 or 12 times-passaged LRC-BCG (LRC-BCG (6) or LRC-BCG (12)). The samples were subjected to western blotting, and PEST-HSP70-MMPII-PEST recombinant protein was detected by αHSP70 (Left) or αMMPII (Right). Arrows show PEST-HSP70-MMPII-PEST protein, and arrowheads show endogenous HSP70

PCR analysis confirmed that LRC-BCG harbors a fusion gene of the expected size integrated into its genome (Fig. 1b). We then examined the expression of the PEST-HSP70-MMPII-PEST recombinant fusion antigen by western blotting. As shown in Figs. 1c and 1d, the recombinant antigen was detected in both the bacterial cells and the culture supernatant of LRC-BCG. Additionally, we examined the genetic stability of LRC-BCG after 12 passages. For practical BCG vaccines, the total number of passages from the master seed to the final lot should not exceed 12^31^. We examined the genomic size of the gene fragment of LRC-BCG which has undergone recombination after 12 passages, and it was as predicted (5023bp) (Fig. 1b). Further, we also detected the PEST-HSP70-MMPII-PEST antigen in both bacterial pellets (Fig. 1c) and culture supernatant (Fig. 1d) derived from LRC-BCG after 12 passages. Based on these results, we successfully generated a novel recombinant BCG expressing the PEST-HSP70-MMPII-PEST antigen by integrating an expression cassette into the BCG-Tokyo genome, indicating the suitability of the recombinant BCG for practical use.

### Macrophage activation by LRC-BCG in vitro

We then compared macrophage activation following infection with BCG-Tokyo and LRC-BCG cells. The murine macrophage cell line J774.1 was infected with the indicated doses of BCG-Tokyo and LRC-BCG for 3 days, and the production of pro-inflammatory cytokines (interleukin [IL]-6, IL-12p40, tumor necrosis factor alpha [TNF-α], and IL-1β) was measured (Fig. 2a). LRC-BCG elicited higher levels of these cytokines than BCG-Tokyo did. We also analyzed inducible nitric oxide synthase (iNOS) mRNA expression using real-time PCR (Fig. 2b) and confirmed that LRC-BCG induced higher iNOS mRNA expression than BCG-Tokyo.

**Fig. 2.**
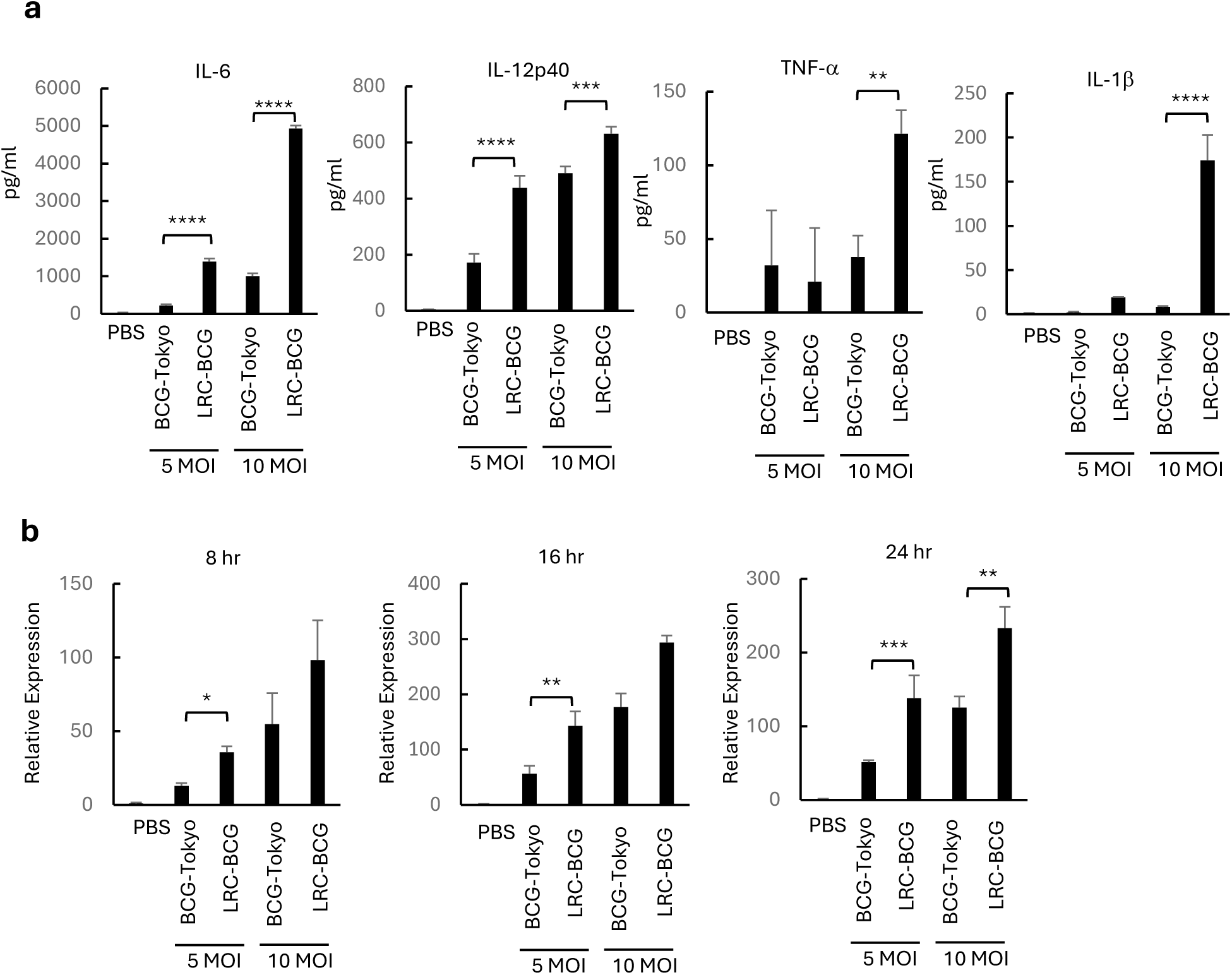
Macrophage activation by LRC-BCG. **a.** Measurement of pro-inflammatory cytokine production from J774.1 macrophage cell line activated with the indicated MOI of BCG-Tokyo or LRC-BCG. **b.** Relative expression of iNOS gene mRNA from J774.1 macrophage cell line activated with the indicated MOI of BCG-Tokyo or LRC-BCG. The expression was normalized by comparing with *Actb* gene expression and data are presented as values relative to the PBS group set as 1. A representative of three separate experiments is shown. Assays were performed in triplicate, and the results are expressed as the mean ± SD. ELISA titers and qPCR ΔCt values were statistically compared using ANOVA followed by Sidak’s multiple comparison test as a post-hoc analysis (*P < 0.05, **P < 0.01, ***P < 0.001, ****P < 0.0001).

### DC activation by LRC-BCG in vitro

We investigated the activation of murine bone marrow-derived dendritic cells (BMDCs) after infection with BCG-Tokyo or LRC-BCG. BMDCs were infected with the indicated dose of BCG-Tokyo and LRC-BCG for 3 days, and production of pro-inflammatory cytokines (IL-6, IL-12p40, and TNF-α) was measured (Fig. 3a).

**Fig. 3.**
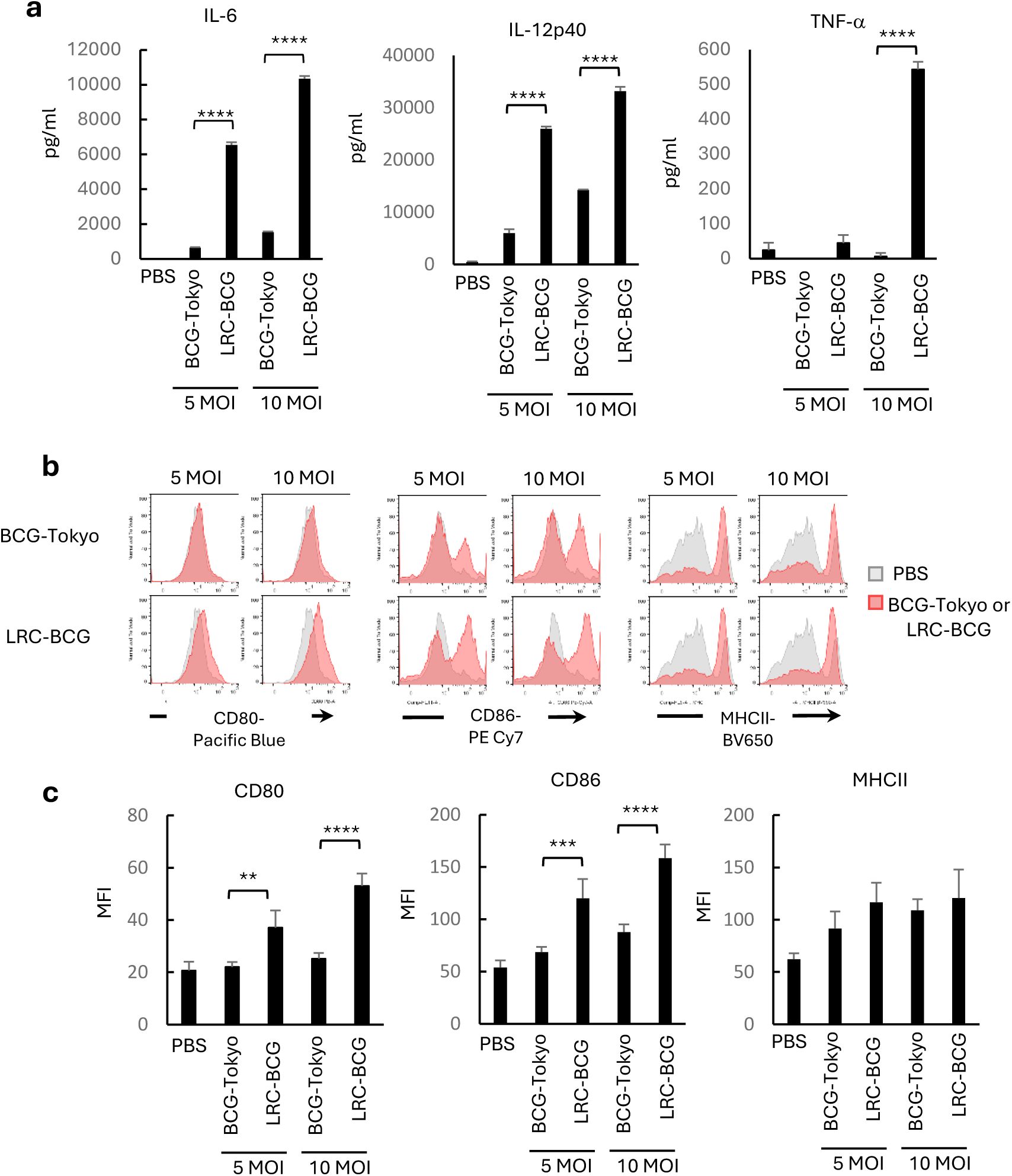
DC activation by LRC-BCG. **a.** Measurement of pro-inflammatory cytokine production from BMDCs activated with the indicated MOI of BCG-Tokyo or LRC-BCG. **b.** Cell surface expression of CD80, CD86 and MHCII on BMDCs activated with the indicated MOI of BCG-Tokyo or LRC-BCG. The histogram of BCG-Tokyo and LRC-BCG (pink) are overlaid with that of PBS group (gray). **c.** Summary of MFI of CD80, CD86 and MHCII on BMDCs shown in **b**. Panels show a representative result from three independent experiments; assays were performed in triplicate within each experiment, and results are expressed as the mean ± SD. Titers were statistically compared using ANOVA followed by Sidak’s multiple comparison test as a post-hoc analysis (**P < 0.01, ***P < 0.001, ****P < 0.001).

We also analyzed the surface expression of CD80, CD86, and MHC class II (MHC II) on BMDCs. The surface expression of CD80 and CD86 was markedly upregulated following infection with LRC-BCG compared to that with BCG-Tokyo (Figs. 3b and 3c). Regarding MHC II upregulation, LRC-BCG showed slightly higher induction than BCG-Tokyo, although the difference was not significant (Figs. 3b and 3c). These results demonstrate that LRC-BCG induces stronger pro-inflammatory activation of macrophages and DCs *in vitro* than BCG-Tokyo.

Together, these findings show that LRC-BCG consistently induces stronger pro-inflammatory activation of both macrophages and BMDCs in vitro than BCG-Tokyo, as reflected in cytokine production and costimulatory molecule upregulation.

### Vaccine efficacy of LRC-BCG against TB in vivo

Our results demonstrate that LRC-BCG activates APCs *in vitro.* We next examined whether these in vitro immunostimulatory properties would translate into protective efficacy *in vivo* (Fig. 4). C57BL/6 mice vaccinated with BCG-Tokyo or LRC-BCG for 4 weeks were challenged with 500 colony-forming units (CFUs) of the MTB H37Rv strain per lung via aerosol. Four weeks later, the mice were sacrificed, and TB in the lungs and spleen was enumerated using the CFU assay. Vaccination with LRC-BCG inhibited the multiplication of MTB in the lungs and the spleens more efficiently than that with BCG-Tokyo (Fig. 4a). Notably, LRC-BCG demonstrated significant protective efficacy even at a low vaccination dose (10³ CFUs/head). The weight of the left lung was measured, and lung tissue samples were analyzed by hematoxylin and eosin (H&E) and Ziehl–Neelsen staining (Figs. 4b-d). Representative H&E-stained lung sections suggested a trend toward reduced inflammation and attenuated pathological changes in the LRC-BCG-inoculated group compared with the BCG-Tokyo group, although this observation was based on qualitative visual inspection rather than quantitative scoring (Fig. 4b). The left lung was visibly heavier in mice with higher bacterial burden, consistent with more severe pulmonary pathology (Fig. 4c). Ziehl–Neelsen staining consistently showed that fewer MTB cells were detected in the lungs of the LRC-BCG-inoculated group than in the BCG-Tokyo group (Fig. 4d). Collectively, these findings provide evidence that LRC-BCG outperformed BCG-Tokyo as a vaccine against MTB in a murine model.

**Fig. 4.**
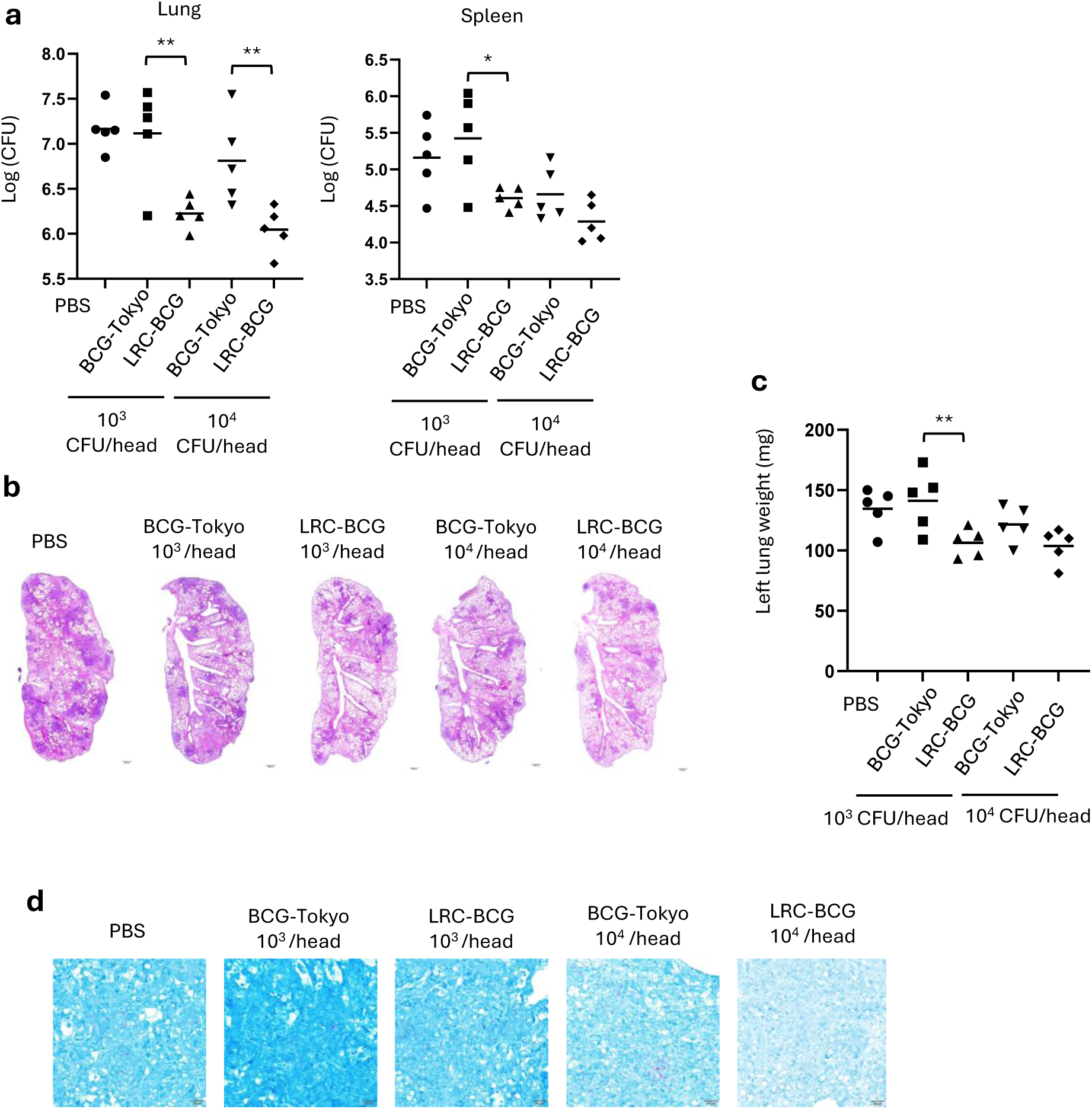
Vaccine efficacy of LRC-BCG. **a.** CFU assay of MTB recovered from lung (left panel) or spleen (right panel). Mice were inoculated with PBS, BCG, or LRC-BCG at 10^3^ or 10^4^ CFUs/head and challenged with MTB H37Rv. MTB recovered from lung and spleen were enumerated by CFU assay. **b.** H&E staining of mouse lung tissue. Representative samples from each group are shown. **c.** Weight of left lungs. **d.** Ziehl-Neelsen staining of mouse lung tissue. Representative samples from each group are shown. Vaccine efficacy was analyzed independently twice, and a representative of the experiments is shown. CFU and lung weight were statistically compared using ANOVA followed by Sidak’s multiple comparison test as a post-hoc analysis (*P < 0.05, **P < 0.01).

## Discussion

In the present study, we identified P79 as a mycobacteriophage-derived promoter that was suitable for genome-integrated antigen expression in BCG. Using this promoter, we successfully generated LRC-BCG, which maintained stable antigen expression and exhibited enhanced immunostimulatory activity and protective efficacy against MTB.

LRC-BCG maintained stable expression of the exogenous antigen across serial passages, a prerequisite for the practical development of vaccines. Functionally, LRC-BCG induced enhanced activation of macrophages and DCs *in vitro* and conferred improved protection against pulmonary TB compared to conventional BCG-Tokyo. Notably, LRC-BCG maintained vaccine efficacy even at low doses.

The successful generation of LRC-BCG suggests that P79 can support recombinant BCG vaccine construction while maintaining stable antigen expression following genome integration. This single-copy, high-level expression capability addresses a practical limitation that has constrained genome-integrated recombinant BCG platforms: genome integration typically results in reduced antigen expression compared to multicopy plasmid systems, owing to the lower gene dosage associated with single-copy insertion. By employing P79, LRC-BCG achieved substantially higher antigen expression than that observed with the conventional Phsp60 promoter in a single-copy context, approaching the expression levels associated with multicopy plasmid-based systems, while avoiding the instability inherent to episomal vectors. A previous study engineered a mycobacteriophage BPs-derived promoter for enhanced activity in an integrated genomic context and proposed its potential utility for high-level antigen expression in recombinant vaccines; however, this work characterized promoter activity using a reporter gene in *Mycobacterium smegmatis* and MTB, and was not directed toward the development of an antigen-expressing recombinant BCG vaccine or the evaluation of protective efficacy *in vivo*^32^.To our knowledge, this is the first report applying a mycobacteriophage-derived promoter to achieve single-copy antigen expression in a genome-integrated recombinant BCG vaccine strain, with demonstrated protective efficacy against TB *in vivo*. In this context, P79 functions as an enabling technology that facilitates the practical implementation of genome-integrated recombinant BCG vaccines. Many studies have generated recombinant BCG expressing exogenous antigens to develop novel, effective vaccines^33,34^. To advance novel recombinant BCG for practical applications, genome-integrated DNA recombination is essential to ensure long-term genetic stability.

LRC-BCG was designed to engage multiple steps of antigen processing and presentation—rather than a single pathway—through *ureC* deletion, antigen secretion, and PEST-mediated degradation, providing a rationale for its enhanced immunostimulatory profile observed *in vitro.* The *in vitro* data indicate that LRC-BCG simultaneously enhances multiple aspects of APC function, including pro-inflammatory activation and upregulation of co-stimulatory molecules. Although these findings do not clarify the causal mechanisms, they are consistent with the notion that LRC-BCG promotes a coordinated immune environment conducive to effective vaccine-induced responses. While Th1-type immunity is widely regarded as a crucial component of host resistance to TB, previous studies have shown that the magnitude of Th1 cytokine responses is not necessarily correlated with vaccine efficacy^35–37^. Further mechanistic studies, including assessment of antigen-specific T-cell responses, will be needed to define the immune correlates underlying LRC-BCG-induced protection.

LRC-BCG conferred greater protection against pulmonary TB than conventional BCG-Tokyo in a murine aerosol challenge model. Novel TB vaccine candidates are generally expected to reduce lung bacterial burden by at least 0.7 log10 CFU relative to unvaccinated controls, based on the level of protection typically observed with BCG vaccination in C57BL/6 mice^38^. In the present study, LRC-BCG vaccination reduced lung bacterial burden by 0.94–1.12 log10 relative to PBS-treated controls across both doses tested. Notably, LRC-BCG exhibited vaccine efficacy even at a low inoculation dose, suggesting a potential dose-sparing effect. Such properties may be advantageous for minimizing vaccine-associated adverse events^39,40^ while maintaining protective capacity.

Our study demonstrates that P79 can be successfully applied to generate a stable genome-integrated recombinant BCG vaccine. The resulting strain, LRC-BCG, showed enhanced protection against tuberculosis in a murine model and represents a promising vaccine candidate for further development. Successful implementation of the LRC-BCG may help reduce the future economic burden on healthcare systems and support progress toward the WHO’s End TB Strategy goals.

## Methods

### Mice

C57BL/6 male mice were purchased from CLEA Japan, Inc. (Tokyo, Japan) and kept in the animal facility of the National Institute of Infectious Diseases. All animals were kept under specific pathogen-free conditions and provided with sterilized food and water.

Mice were euthanized under isoflurane anesthesia. The Animal Research Committee of Experimental Animals at the National Institute of Infectious Diseases reviewed and approved all animal studies conducted in accordance with their guidelines.

### Bacterial culture

BCG-Tokyo, LRC-BCG, and MTB H37Rv strain were cultured at 37 °C in 7H9 medium supplemented with Tween 80 and Middlebrook ADC Enrichment (BD Biosciences, Franklin Lakes, NJ, USA) and subcultured twice a week. All bacteria were harvested at the log phase, aliquoted, and stored at -80 °C. Frozen stocks were thawed and counted using a CFU assay before use. For the in vivo comparison of vaccine efficacy, we used commercial BCG (Japan BCG Laboratory, Tokyo, Japan) as the control BCG-Tokyo to evaluate vaccine efficacy under conditions more relevant to practical use.

### Analysis of transcriptional activity

P79 promoter was cloned from mycobacteriophage TM4 gDNA using the primers F P79 Xba/R P79 Eco. EGFP was amplified with the primers F15P79 EG/R 15ter1 EG 780.

The sequences of the primers are provided in Supplementary Table 1. The P79 promoter and EGFP were ligated and then inserted into pMV261 and pMV306. Mock vectors and EGFP-expressing plasmids were introduced into BCG-Tokyo via electroporation. The transformed BCGs were selected by adding kanamycin to the culture plates.

Recombinant BCGs containing plasmids were isolated, and EGFP expression was quantified using BD FACS Canto II (BD Biosciences). The data were analyzed using FlowJo v10 software (BD Biosciences).

### LRC-BCG generation

As illustrated in Supplementary Fig. 2, the donor DNA fragment was constructed as follows. First, the P79-EGFP-pMV306 plasmid was digested with EcoRI and HpaI and assembled with PEST-HSP70-MMPII-PEST amplified using primers F 15 P79 70B / R 15ter MMP, with the In-Fusion HD Cloning Kit (Takara Bio, Shiga, Japan). Then the expression cassette including P79, PEST-HSP70-MMPII-PEST, and ter sequence was amplified with primers F P79XbaI/ R ter SmaI. Second, the upstream region (UR UP) and downstream region (UR DO) of the *ureC* gene, amplified from BCG-Tokyo gDNA using primers F UreUP HindIII/R UreUP XbaI and F UreDO SmaI/R UreDO EcoRI respectively, were ligated to the expression cassette via restriction enzyme digestion into pUC18. Subsequently, the SmaI site located between the ter sequence and UR DO was digested, and the Hyg^r^ was inserted. The resulting plasmid consisted of UR UP, P79, PEST-HSP70-MMPII-PEST, ter sequence, Hyg^r^, and UR DO. Finally, the donor DNA fragment for generating LRC-BCG (Fig. 1a) was prepared by amplifying with the primers M4-2 and RV-2. The sequences of the primers are shown in Supplementary Table 1.

Recombinant BCG was generated using a mycobacterial phage-recombinase system^23,41–43^. Briefly, BCG-Tokyo cells were electroporated with the pJV53 plasmid and selected for kanamycin resistance. BCG-Tokyo cells transformed with pJV53 were then electroporated with the donor DNA fragment, and recombinant BCG was selected for hygromycin resistance. To remove Hyg^r^, the pYUB870 plasmid carrying a res site-specific resolvase was introduced into the recombinant BCG. After kanamycin selection, the absence of Hyg^r^ was confirmed by PCR using primers UreC-1211 and UreC+1703. Hyg^r^-free recombinants were then cultured in 7H9 broth without antibiotics, and pYUB870-free recombinant BCG was selected to yield LRC-BCG.

### Western blotting

BCG-Tokyo and LRC-BCG were cultured, and the bacterial pellets were collected to confirm the expression of PEST-HSP70-MMPII-PEST, an exogenous antigen derived from LRC-BCG. The collected pellet was suspended in PBS and sonicated for 10 min using a Bioruptor (Sonicbio, Kanagawa, Japan). The sonicated samples were centrifuged, and the supernatants were subjected to western blotting. To verify the secretion of exogenous antigens, BCG-Tokyo, and LRC-BCG were cultured for 14 days in Sauton’s medium, and the culture supernatants were collected. The supernatant was concentrated using an Amicon Ultra (Merck Millipore, Darmstadt, Germany) after BCG depletion by centrifugation. SDS-PAGE and electroblotting were conducted using standard methods. Western blotting was performed as follows: the membrane was blocked with Block Ace Solution (KAC Co., Ltd., Kyoto, Japan) and incubated with anti-MMPII mAb 202-3 or anti-mycobacterial HSP70 mAb (HyTEST, Turku, Finland). Alkaline phosphatase-conjugated anti-mouse IgG antibody (Jackson Immunoresearch, West Grove, PA, USA) was used as the secondary antibody. Color development was performed using NBT/5-bromo-4-chloro-3-indolyl phosphate detection reagent (Calbiochem, San Diego, CA, USA).

### In vitro culture of the J774.1 cell line and murine BMDCs

The J774.1 cell line was cultured and passaged in RPMI 1640 medium (Thermo Fisher Scientific, Waltham, MA, USA) supplemented with 10% FCS, 2-mercaptoethanol, and penicillin G. J774.1 was cultured at a density of 1 × 10^5^ cells/200 μL/well in a 96-well culture plate (Corning, Corning, NY, USA) and infected with BCG-Tokyo or LRC-BCG at the indicated multiplicity of infection.

For the generation of BMDCs, bone marrow (BM) cells were collected from the tibiae, femora, and pelvises of mice. BM cells were then plated at a density of 1 × 10^6^ cells/mL/well in a 24-well culture plate (Corning) and cultured in complete RPMI 1640 culture medium supplemented with 10 ng/mL recombinant murine GM-CSF (Thermo Fisher Scientific). The cultures were fed every 2 days by gently swirling the plates, aspirating 50% of the medium, and replenishing it with fresh medium supplemented with GM-CSF. BMDCs were collected on day 7, plated at a density of 1 × 10^5^ cells/200 μL/well in 96-well culture plates (Corning), and infected with BCGs.

### ELISA assay

IL-1β, IL-6, IL-12p40, and TNF-α production from J774.1 and BMDC were measured by ELISA Assay using OptEIA ELISA kit (BD Biosciences) following the manufacturer’s instructions.

### Real-time PCR analysis of mRNA expression

J774.1 was harvested after the indicated time of infection with BCG-Tokyo or LRC-BCG. Cells were lysed, and total RNA was purified using a Qiagen Miniprep kit (Qiagen, Germantown, MD, USA). Complementary DNA (cDNA) was synthesized from RNA using the ReverTra Ace qPCR RT Master Mix with gDNA Remover (Toyobo, Osaka, Japan). The primers used for real-time PCR analysis were purchased from Takara Bio. The nucleotide sequences of the primers were as follows: Actb forward primer: 5′-CATCCGTAAAGACCTCTATGCCAAC-3′, Actb reverse primer: 5′-ATGGAGCCACCGATCCACA-3′, Nos2 forward primer: 5′-ATGGCTCGGGATGTGGCTAC-3′, Nos2 reverse primer: 5′-AAAGACTGCACCGAAGATATCTTCA-3′. cDNA was amplified using the primers and THUNDERBIRD Next SYBR qPCR Mix (Toyobo), and amplification was measured using QuantStudio Pro 7 (Thermo Fisher Scientific). Actb served as the reference gene, and group differences were assessed using the ΔCt values for each sample.

### Flow cytometry analysis

BMDCs were collected, and Fc receptors on the cells were blocked using FcX Plus (BioLegend, San Diego, CA, USA). To exclude dead cells from the analysis, cells were stained with Zombie Aqua™ Fixable Viability Kit (BioLegend). The cells were stained with the antibodies below, fixed in 4% paraformaldehyde, and analyzed using a MACSQuant Analyzer 16 Flow Cytometer (Miltenyi Biotec, Bergisch Gladbach, Germany). Data were analyzed using FlowJo software (BD). The antibodies used in this study were αCD80-Pacific Blue (clone 16-10A1), αCD86-PECy7 (Clone: PO3), αCD11c-Brilliant Violet 570 (Clone: N418), and αMHCII-Brilliant Violet 650 (Clone: M5/114.15.2). All antibodies were purchased from BioLegend.

### Vaccine efficacy analysis in mice

To observe the effect of BCG vaccination on MTB infection, five C57BL/6 mice per group were inoculated with PBS, BCG-Tokyo or LRC-BCG at the indicated doses. Four weeks later, the mice were challenged with 500 CFUs/lung of the H37Rv strain via aerosol infection using an automated inhalation exposure apparatus (Glas-Col Corp., IN, Model 099C A4212). Four weeks later, the challenged mice were sacrificed, and the lungs and the spleens were carefully excised and collected. The right lungs and the spleens were mechanically disrupted using Gentle MACS (Miltenyi Biotec), and the bacterial burden was enumerated using a colony assay. The left lung was weighed and then fixed for histopathological analysis (Applied Medical Research Laboratory, Osaka, Japan).

### Statistics

Lung and spleen CFU counts were log10-transformed prior to statistical analysis. For multiple comparisons, one-way analysis of variance (ANOVA) followed by Sidak’s multiple comparison test as a post-hoc analysis was employed with GraphPad Prism 9 (GraphPad Software, Boston, MA, USA). All tests were two-tailed, and statistical significance was set as P < 0.05.

## Data availability

All data supporting the findings of this study are available from the corresponding author upon reasonable request.

## Code availability

Not applicable.

## Acknowledgements

This research was supported by AMED under Grant Numbers JP22fk0108648 to Y. Tsujimura, JP16fk0108105, and JP19fk0108089 to T. T., and was partially supported by JP223fa727002 to M. A. We thank Ms. M. Kujiraoka for her technical support, and Dr. Saburo Yamamoto and Dr. Toshiko Yamamoto for their helpful advice and comments on this study. We would also like to thank Editage (www.editage.jp) for English language editing.

## Author Contributions

Y. Tsukamoto and T. T. designed the project. Y. Tsukamoto, T. T., M. M. and M. A. wrote the manuscript. T. M., Y. Maeda, and Y. Miyamoto generated and prepared the BCGs. Y. Tsukamoto, Y. Tsujimura, S. T., H. S., M. N. A., T. K., and M. N. conducted the experiments and analyzed the data. All authors have reviewed and approved the final manuscript.

## References

1. 1. WHO. Global Tuberculosis Report 2025. (WHO, 2025).

2. Pottenger, F. M. Tuberculosis: Vaccination Against Tuberculosis with B. C. G. (Calmette). Cal West Med 30, 131–2 (1929).

3. Lange, C. et al. 100 years of Mycobacterium bovis bacille Calmette-Guerin. Lancet Infect Dis 22, e2–e12 (2022).

4. Migliori, G. B. et al. History of prevention, diagnosis, treatment and rehabilitation of pulmonary sequelae of tuberculosis. Presse Med 51, 104112 (2022).

5. 5. WHO. BCG Vaccines: WHO Position Paper – February 2018. (WHO, 2018).

6. Trunz, B. B., Fine, P. & Dye, C. Effect of BCG vaccination on childhood tuberculous meningitis and miliary tuberculosis worldwide: a meta-analysis and assessment of cost-effectiveness. Lancet 367, 1173–80 (2006).

7. Abubakar, I. et al. Systematic review and meta-analysis of the current evidence on the duration of protection by bacillus Calmette-Guerin vaccination against tuberculosis. Health Technol Assess 17, 1–372, v–vi (2013).

8. Roy, A. et al. Effect of BCG vaccination against Mycobacterium tuberculosis infection in children: systematic review and meta-analysis. BMJ 349, g4643 (2014).

9. Fine, P. E. Variation in protection by BCG: implications of and for heterologous immunity. Lancet 346, 1339–45 (1995).

10. Martinez, L. et al. Infant BCG vaccination and risk of pulmonary and extrapulmonary tuberculosis throughout the life course: a systematic review and individual participant data meta-analysis. Lancet Glob Health 10, e1307–e1316 (2022).

11. Kaufmann, S. H. E. The Paediatric BCG Vaccine Century: From Historical Success to Future Innovations. Acta Paediatr https://doi.org/10.1111/apa.70182 (2025) doi:10.1111/apa.70182.

12. Elbehiry, A. et al. Advancing the fight against tuberculosis: integrating innovation and public health in diagnosis, treatment, vaccine development, and implementation science. Front Med Lausanne 12, 1596579 (2025).

13. Konjengbam, B. D., Meitei, H. N., Pandey, A. & Haobam, R. Goals and strategies in vaccine development against tuberculosis. Mol Immunol 183, 56–71 (2025).

14. Maeda, Y., Mukai, T., Spencer, J. & Makino, M. Identification of an Immunomodulating Agent from Mycobacterium leprae. Infect Immun 73, 2744–50 (2005).

15. Makino, M., Maeda, Y. & Ishii, N. Immunostimulatory activity of major membrane protein-II from Mycobacterium leprae. Cell Immunol 233, 53–60 (2005).

16. Makino, M., Maeda, Y. & Inagaki, K. Immunostimulatory activity of recombinant Mycobacterium bovis BCG that secretes major membrane protein II of Mycobacterium leprae. Infect Immun 74, 6264–71 (2006).

17. Tsukamoto, Y. et al. Immunostimulatory activity of major membrane protein II from Mycobacterium tuberculosis. Clin Vaccine Immunol 18, 235–42 (2011).

18. Mukai, T., Tsukamoto, Y., Maeda, Y., Tamura, T. & Makino, M. Efficient activation of human T cells of both CD4 and CD8 subsets by urease-deficient recombinant Mycobacterium bovis BCG that produced a heat shock protein 70-M. tuberculosis-derived major membrane protein II fusion protein. Clin Vaccine Immunol 21, 1–11 (2014).

19. Tsukamoto, Y., Maeda, Y., Tamura, T., Mukai, T. & Makino, M. Polyclonal activation of naive T cells by urease deficient-recombinant BCG that produced protein complex composed of heat shock protein 70, CysO and major membrane protein-II. BMC Infect Dis 14, 179 (2014).

20. Tsukamoto, Y. et al. Enhanced protective efficacy against tuberculosis provided by a recombinant urease deficient BCG expressing heat shock protein 70-major membrane protein-II having PEST sequence. Vaccine 34, 6301–6308 (2016).

21. Miyamoto, Y. et al. Production of antibiotic resistance gene-free urease-deficient recombinant BCG that secretes antigenic protein applicable for practical use in tuberculosis vaccination. Tuberc. Edinb 129, 102105 (2021).

22. Grode, L. et al. Increased vaccine efficacy against tuberculosis of recombinant Mycobacterium bovis bacille Calmette-Guerin mutants that secrete listeriolysin. J Clin Invest 115, 2472–9 (2005).

23. Mukai, T., Maeda, Y., Tamura, T., Miyamoto, Y. & Makino, M. CD4+ T-cell activation by antigen-presenting cells infected with urease-deficient recombinant Mycobacterium bovis bacillus Calmette-Guerin. FEMS Immunol Med Microbiol 53, 96–106 (2008).

24. Liu, P., Sang, Z., Liu, K., Zhang, M. & Niu, Y. Mycobacterium tuberculosis Hsp70 as a cancer vaccine adjuvant: Immunomodulatory mechanisms and tumor microenvironment remodeling. Vaccine 62, 127493 (2025).

25. Bulut, Y. et al. Mycobacterium tuberculosis heat shock proteins use diverse Toll-like receptor pathways to activate pro-inflammatory signals. J Biol Chem 280, 20961–7 (2005).

26. Tucci, P., Portela, M., Chetto, C. R., Gonzalez-Sapienza, G. & Marin, M. Integrative proteomic and glycoproteomic profiling of Mycobacterium tuberculosis culture filtrate. PLoS One 15, e0221837 (2020).

27. Rechsteiner, M. & Rogers, S. W. PEST sequences and regulation by proteolysis. Trends Biochem Sci 21, 267–71 (1996).

28. Decatur, A. L. & Portnoy, D. A. A PEST-like sequence in listeriolysin O essential for Listeria monocytogenes pathogenicity. Science 290, 992–5 (2000).

29. Kaufmann, S. H. Tuberculosis vaccine development: strength lies in tenacity. Trends Immunol 33, 373–9 (2012).

30. Stover, C. K. et al. New use of BCG for recombinant vaccines. Nature 351, 456–60 (1991).

31. 31. WHO. Requirements for Freeze-Dried BCG Vaccine-WHO Technical Report Series. vol. 745 (WHO Expert Committee on Biological Standardization, 1987).

32. Oldfield, L. M. & Hatfull, G. F. Mutational Analysis of the Mycobacteriophage BPs Promoter PR Reveals Context-Dependent Sequences for Mycobacterial Gene Expression. J. Bacteriol. 196, 3589–3597 (2014).

33. Mouhoub, E., Domenech, P., Ndao, M. & Reed, M. B. The Diverse Applications of Recombinant BCG-Based Vaccines to Target Infectious Diseases Other Than Tuberculosis: An Overview. Front. Microbiol. 12, 757858 (2021).

34. Nieuwenhuizen, N. E. & Kaufmann, S. H. E. Next-Generation Vaccines Based on Bacille Calmette–Guérin. Front. Immunol. 9, 121 (2018).

35. North, R. J. & Jung, Y. J. Immunity to tuberculosis. Annu Rev Immunol 22, 599– 623 (2004).

36. Salgame, P. Host innate and Th1 responses and the bacterial factors that control Mycobacterium tuberculosis infection. Curr Opin Immunol 17, 374–80 (2005).

37. Zeng, G., Zhang, G. & Chen, X. Th1 cytokines, true functional signatures for protective immunity against TB? Cell Mol Immunol 15, 206–215 (2018).

38. Singh, A. K. & Gupta, U. D. Animal models of tuberculosis: Lesson learnt. Indian J. Med. Res. 147, 456–463 (2018).

39. Horikoshi, Y. & Toizumi, M. Pediatric tuberculosis and BCG vaccine in Japan. Vaccine 62, 127564 (2025).

40. Okuno, H. et al. Characteristics and incidence of vaccine adverse events after Bacille Calmette-Guerin vaccination: A national surveillance study in Japan from 2013 to 2017. Vaccine 40, 4922–4928 (2022).

41. Snapper, S. B., Melton, R. E., Mustafa, S., Kieser, T. & Jacobs, W. R., Jr. Isolation and characterization of efficient plasmid transformation mutants of Mycobacterium smegmatis. Mol Microbiol 4, 1911–9 (1990).

42. van Kessel, J. C. & Hatfull, G. F. Recombineering in Mycobacterium tuberculosis. Nat Methods 4, 147–52 (2007).

43. Kawakita, T. et al. Point mutation in the stop codon of MAV_RS14660 increases the growth rate of Mycobacterium avium subspecies hominissuis. Microbiology 167, 001007 (2021).

