## Supplementary Materials for "A novel recombinant BCG vaccine using a mycobacteriophage promoter shows improved protection against tuberculosis"

### Supplementary Fig. 1

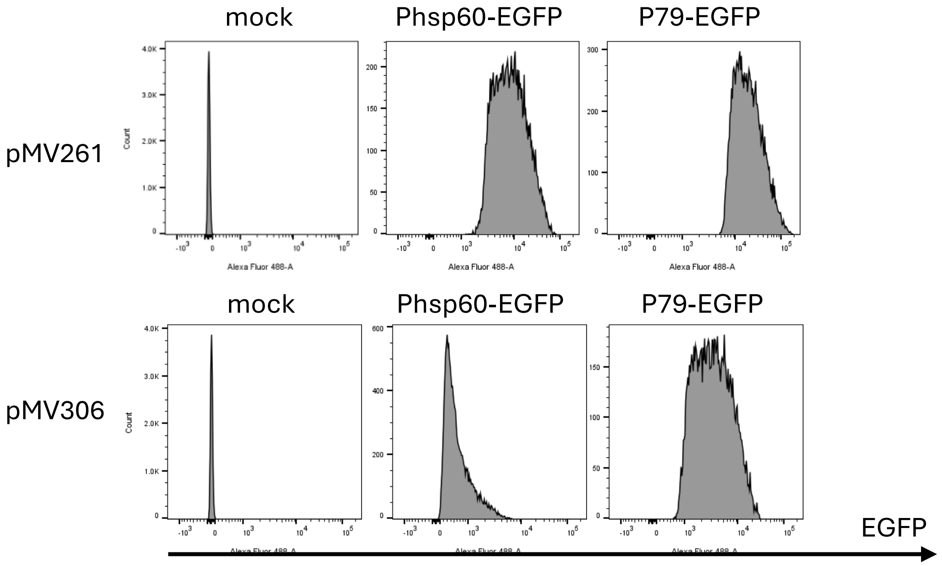


**Supplementary Fig. 1** Comparison of transcriptional activity between promoters Phsp60 and P79
BCG-Tokyo was introduced with pMV261-based plasmid (above) or pMV306-based plasmid (below) and intensity of EGFP was measured by flow cytometry. BCG was introduced with mock vector (left), plasmids containing EGFP downstream of promoter Phsp60 (center) or plasmids containing EGFP downstream of P79 (right).

Supplementary Fig. 2

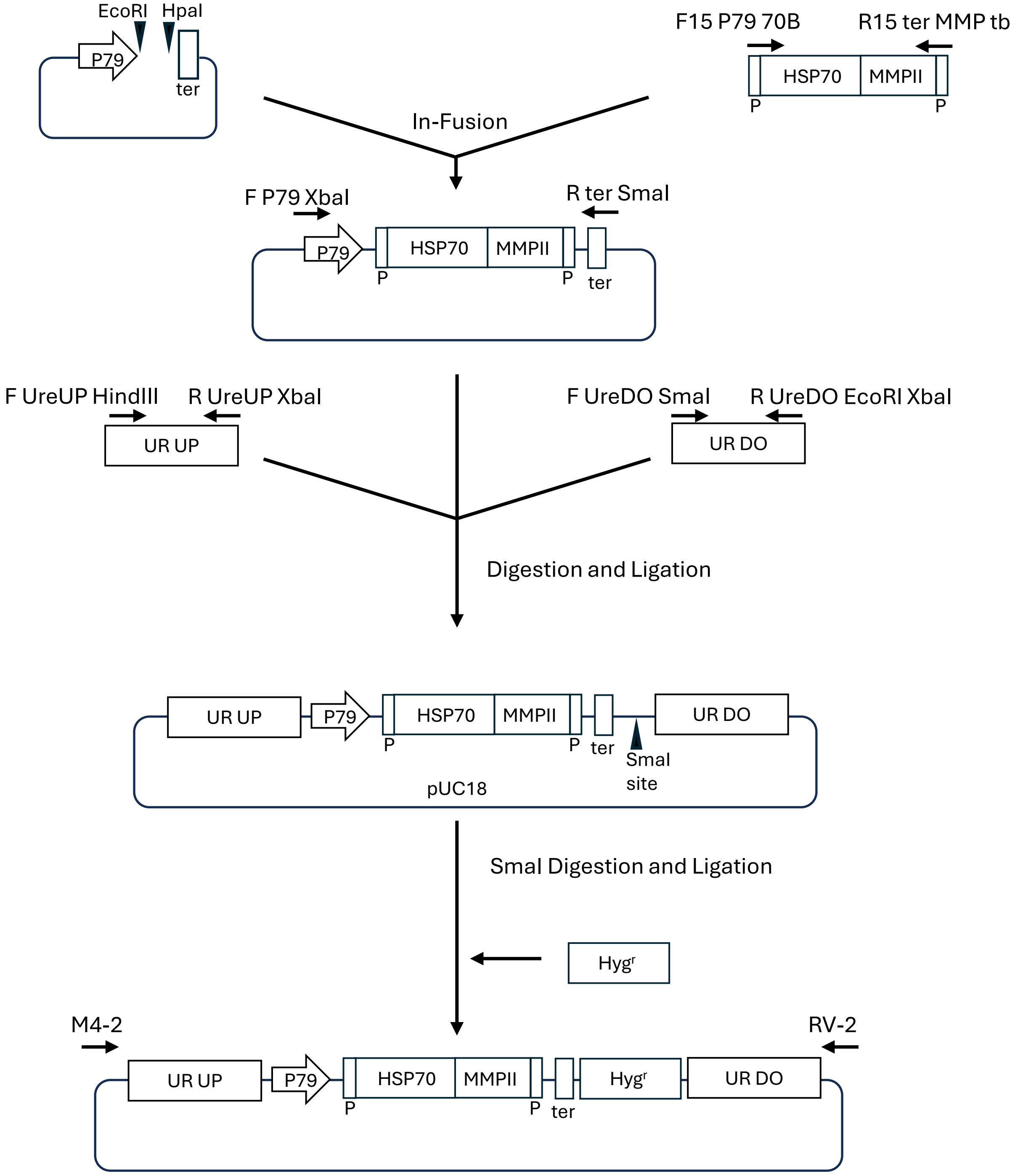


**Supplementary Fig. 2** Construction of plasmids for generating donor DNA fragment Schematic diagram of donor DNA fragment generation. P and ter indicate PEST sequence and transcription terminator. UR UP and UR DO indicate the upstream and downstream regions of the *ureC* gene respectively (see Methods). UR UP, P79 promoter, PEST-HSP70-MMPII-PEST, ter, Hyg$^r$ and UR DO were inserted into pUC18 vector and amplified by M4-2 and RV-2 primers to generate donor DNA fragment.

**Supplementary Table 1.** Sequence of primers used to generate recombinant BCGs

| Primer | Sequence (5′ to 3′) |
|---|---|
| F P79 Xba | TTCTCTAGACACCGCATCAACAAAACCC |
| R P79 Eco | ATCGAATTCGAGGCCCTCCTCGGGCT |
| F15P79 EG | CCCGAGGAGGGCCTCATGGTGAGCAAGGGCGAGG |
| R 15ter1 EG 780 | ATGCCTAGTTAACTACTTGTACAGCTCGTCCAT |
| F15 P79 70B | CCCGAGGAGGGCCTCATGGCTCGTGCGGTCGG |
| R15 ter MMP | TACGCTAGTTAACTATCAGGTCGGTGGGCGAGA |
| F P79XbaI | TTCTCTAGACACCGCATCAACAAAACCC |
| R ter SmaI | AATTACCCGGGTGATCACCGCGGCCATGAT |
| F UreUP HindIII | ACACAAGCTTGTTGGTGTAGACACAAGGAC |
| R UreUP XbaI | TTTTTCTAGAAACCCGACGATTTGGGGAAT |
| F UreDO SmaI | TTTTCCCGGGATACGGTGAACACCCTTGAC |
| R UreDO EcoRI | GTGTGAATTCTTCAGGAACGCTTCCAGGGT |
| M4-2 | TCTTCGCTATTACGCCAGCT |
| RV-2 | GTTGTGTGGAATTGTGAGCG |
| F UreC-1211 | AAATACCGTTGTGTCATAGGT |
| R UreC+1703 | TCATTACTTCGTCGAGCCAC |
